# Maternal Overprotection in Childhood is Associated with Amygdala Reactivity and Structural Connectivity in Adulthood

**DOI:** 10.1101/535823

**Authors:** Madeline J. Farber, M. Justin Kim, Annchen R. Knodt, Ahmad R. Hariri

## Abstract

Recently, we reported that variability in early-life caregiving experiences maps onto individual differences in threat-related brain function. Here, we extend this work to provide further evidence that subtle variability in specific features of early caregiving shapes structural and functional connectivity between the amygdala and medial prefrontal cortex (mPFC) in a cohort of 312 young adult volunteers. Multiple regression analyses revealed that participants who reported higher maternal overprotection exhibited increased amygdala reactivity to explicit signals of interpersonal threat but not implicit signals of broad environmental threat. While amygdala functional connectivity with regulatory regions of the mPFC was not significantly associated with maternal overprotection, participants who reported higher maternal overprotection exhibited relatively decreased structural integrity of the uncinate fasciculus (UF), a white matter tract connecting these same brain regions. There were no significant associations between structural or functional brain measures and either maternal or paternal care ratings. These findings suggest that an overprotective maternal parenting style during childhood is associated with later functional and structural alterations of brain regions involved in generating and regulating responses to threat.

## INTRODUCTION

A rich literature details the widespread effects of early caregiving on child psychosocial development (Belsky & Haan, 2011; Callaghan & Tottenham, 2016; Tottenham, 2018). Seminal early work revealed the formative and lasting impacts of parenting style and attachment on socioemotional development over time across species (Ainsworth, 1969; Bowlby, 1958; Harlow, 1961; Harlow & Zimmermann, 1959; Lorenz, 1935). As a natural extension of these findings, there is a large body of research linking stressful early environments, such as those marked by trauma, abuse, and neglect, with similar outcomes in humans. Children exposed to early life adversity have poorer outcomes in terms of social, emotional, and behavioral development (Gee, 2016; Gee & Casey, 2015; Green et al., 2010; McLaughlin et al., 2010; McLaughlin, Peverill, Gold, Alves, & Sheridan, 2015; Tottenham, 2013). Such neuroimaging research has described parallel effects in brain with children exposed to trauma, abuse, and neglect in early life exhibiting maladaptive alterations in the structure and function of brain regions supporting emotional behaviors, particularly the amygdala and regulatory regions of the medial prefrontal cortex (mPFC).

Taken together, the above research suggests that many of the effects of early caregiving may be determined by the shaping of brain circuits supporting emotional behaviors and psychological well-being (Belsky & Haan, 2011; Callaghan & Tottenham, 2016; Gee, 2016; Tottenham, 2018). However, the majority of this research has focused nearly exclusively on extremes of early life adversity such as trauma, abuse, neglect, and institutionalization (Gee et al., 2013; Herringa et al., 2013; Malter Cohen, Tottenham, & Casey, 2013; McLaughlin et al., 2015; Pechtel, Lyons-Ruth, Anderson, & Teicher, 2014; Sheridan, Fox, Zeanah, McLaughlin, & Nelson, 2012; Tottenham, 2012; Tottenham et al., 2010). With few exceptions (Farber et al., 2018; Romund et al., 2016; Taylor, Eisenberger, Saxbe, Lehman, & Lieberman, 2006; Whittle et al., 2014, 2016, 2009, 2012), there is little work to date investigating potential impacts of normative variability in early caregiving on the development of these brain circuits in the absence of such harsh early life environments.

Recently, we examined associations between variability in caregiving and threat-related amygdala reactivity in a cohort of 232 adolescents (Farber et al., 2018). In this work, we modeled the childhood caregiving environment using the general functioning and affective responsiveness scales of the Family Assessment Device (FAD). Our analyses revealed that greater familial affective responsiveness (i.e., the appropriate expression and recognition of emotion through warmth, care, and affection) was associated with increased amygdala reactivity to explicit, interpersonal threat—as conveyed by angry facial expressions—but not implicit, environmental threat—as conveyed by fearful facial expressions. This association was robust to the potential influence of participant sex, age, broad familial risk for depression, and early life stress, as well as contemporaneous symptoms of depression and anxiety. Moreover, this association was moderated by the experience of recent stressful life events wherein higher affective responsiveness was associated with higher amygdala reactivity in participants reporting low but not high recent stress. In contrast, there were no significant associations between amygdala reactivity and the FAD scale for general family functioning (e.g., “We don’t get along well together”).

This work suggests that adolescents who report less stressful environments outside of the home and home environments marked by greater affective responsiveness exhibit increased amygdala reactivity to interpersonal threat. We hypothesized that this paradoxical association may reflect increased associative learning following less-frequent, more unpredictable experiences of threat or conflict. We further speculated that our observed associations among better familial affective responsiveness, less stress, and higher amygdala reactivity suggest a mechanism through which parental overprotection may manifest as psychosocial dysfunction; however, we had no measure of parental overprotection available in this dataset to test this hypothesis directly.

While this prior work extends the literature on the impact of caregiving extremes on behaviorally-relevant brain function, the data available were not ideal for assessing caregiving in fine detail as the FAD does not provide information on family structure or parent-of-origin effects. Additionally, as previously mentioned, the FAD does not generate indices of the extent to which caregivers were permissive or controlling, two facets of particular importance in shaping the early caregiving environment (Parker, 1983). Parental overprotection, defined as “restrictive and controlling parenting” has been associated with later psychopathology including depression and anxiety disorders (Thomasgard & Metz, 1993). In contrast, families marked by high parental care and low parental overprotection have been described as having “optimal bonding,” with children from such families reporting less distress, better general well-being, and better social support (Canetti, Bachar, Galili-Weisstub, De-Nour, & Shalev, 1997).

In the present study, we sought to extend our prior finding in adolescents and directly test our speculation regarding the role of parental overprotection. We did this by capturing more detailed aspects of early caregiving experiences as well as additional features of corticolimbic circuit integrity using data from 312 young adult volunteers who completed the Duke Neurogenetics Study. First, we expand upon broadband family functioning to parse the specific dimensions of (a) care and (b) control/overprotection—for mothers and fathers independently. Second, we broaden our neuroimaging analyses beyond threat-related amygdala reactivity to also examine both functional and structural connectivity of the amygdala with regulatory regions of the mPFC. Early parenting style was indexed by the Parental Bonding Instrument (PBI; Parker, Tupling, & Brown, 1979) which examines paternal and maternal caregiving separately along the dimensions of care and protection. Throughout our analyses, we controlled for potential confounding effects of (a) early life adversity and (b) socioeconomic status. We utilized this approach given the well documented associations between early life adversity and corticolimbic circuitry (Eluvathingal et al., 2006; Gee et al., 2013; Hanson et al., 2015; Sheridan, Sarsour, Jutte, D’Esposito, & Boyce, 2012), as well as the body of literature maintaining that optimal parenting style varies based on socioeconomic status and related chronic stressors (Roubinov & Boyce, 2017).

As in our prior work, amygdala reactivity to explicit, interpersonal and implicit, environmental threat as communicated by angry and fearful facial expressions, respectively, was measured using task-based functional magnetic resonance imaging (fMRI). Given the complex interpersonal dynamics at play within parent-child interactions and in light of our previous findings, we focused on amygdala reactivity to interpersonal threat cues as specifically indexed by angry facial expressions (Adams, Gordon, Baird, Ambady, & Kleck, 2003). We did, however, test for associations with fearful facial expressions as well to explore specificity of associations. Moreover, we examined several additional features of corticolimbic circuit function and structure. Seed-based amygdala functional connectivity with the mPFC was modeled using generalized psychophysiological interaction (gPPI). The mPFC was targeted because of its important reciprocal connections with the amygdala supporting the integration and regulation of threat-related processing. Accordingly, structural connectivity between these regions was assessed using diffusion weighted imaging-based fractional anisotropy (FA) estimates of white matter microstructural integrity of the uncinate fasciculus (UF), a major structural pathway between the amygdala and mPFC.

Building on our previous work, we hypothesized that higher parental care (both maternal and paternal) experienced during childhood (i.e., before 18 years of age) would be associated with increased amygdala reactivity to angry but not fearful facial expressions in young adulthood (i.e., 18-22 years of age). Extending this primary hypothesis, we explored the following related questions: (1) are maternal and paternal care and overprotection differentially associated with amygdala reactivity, (2) are parental care and overprotection associated with functional connectivity between the amygdala and mPFC, (3) are parental care and overprotection associated with structural connectivity between the amygdala and mPFC, and (4) are potential associations reflective of effects of parenting style above and beyond those of early life adversity and socioeconomic status?

## MATERIALS & METHODS

### Participants

Study participants included a subset of individuals (*N* = 312) having completed the Duke Neurogenetics Study (DNS), which was designed to identify biomarkers of risk for psychopathology amongst young adult university students. The present analyses focus on a substantially smaller subsample of the full DNS sample (*N* = 1332) because the measure of caregiving was added to the DNS protocol during the final year of data collection. All procedures were approved by the Duke University Medical Center Institutional Review Board and all participants provided informed consent before study initiation. Participants in the present DNS subsample (a) were free of cancer, stroke, diabetes, chronic kidney or liver disease, hypertension, or psychotic symptoms; (b) were not actively using psychotropic, glucocorticoid, or hypolipidemic medication; and (c) met quality control for MRI data as described below. In addition to a formal clinical screening for past and current mental illness, all participants provided extensive self-report measures related to behavior and life experiences. All participants further completed a neuroimaging protocol on one of two research-dedicated GE MR750 3T scanners equipped with high-power high-duty-cycle 50-mT/m gradients at 200 T/m/s slew rate, and an eight-channel head coil for parallel imaging at high bandwidth up to 1MHz at the Duke-UNC Brain Imaging and Analysis Center.

As the DNS seeks to examine the broad distribution of dimensional behavioral and biological variables, diagnosis of any past or current DSM-IV Axis I disorder or select Axis II disorders (antisocial personality disorder and borderline personality disorder) was not an exclusion to participation. However, no individuals, regardless of diagnosis, were taking any psychoactive medication during or at least 14 days prior to their participation. Categorical diagnosis was assessed with the electronic Mini International Neuropsychiatric Interview (Lecrubier et al., 1997) and Structured Clinical Interview for the DSM-IV Axis II subtests (First, Gibbon, Spitzer, & Williams, 1996). Of the 312 participants included in our analyses, 14 met criteria for major depressive disorder, 3 for bipolar disorder, 11 for panic disorder and/or agoraphobia, 2 for social anxiety disorder, 3 for generalized anxiety disorder, 45 for alcohol abuse, 12 for substance abuse, 1 for eating disorder(s), and 2 for psychotic symptoms.

### Self-Report Measures

Parental care and overprotection were assessed using the Parental Bonding Instrument (PBI), a 40-item scale used across a variety of research contexts with acceptable validity, reliability, and stability (Murphy, Wickramaratne, & Weissman, 2010; Parker, 1983, 1989; Parker et al., 1979). Of note, the PBI is intentionally retrospective; the measure itself dictates that participants over 16 years of age complete the questionnaire for how they remember their parents during their first 16 years (Parker et al., 1979).

The PBI consists of two separate scales for each parent—care and overprotection—and participants rate the extent to which each item corresponds with the attitudes and behaviors of their parents “when [they] were growing up.” Scores range from 1 (“very like”) to 4 (“very unlike”) with higher scores reflecting higher parental care/overprotection. In addition, the PBI generates separate maternal and paternal quadrants based on the rater’s maternal and paternal care and overprotection scores. The parenting style represented by each quadrant are labeled as “weak” (low care, low overprotection), “affectionless control” (low care, high overprotection), “autonomy support” (high care, low overprotection), and “affective constraint” (high care, high overprotection). However, we focused on dimensional indices of care and overprotection rather than categorical quadrant placements to better model subtle variability in parenting style onto brain continuously rather than categorically.

Early life adversity was assessed using the Childhood Trauma Questionnaire (CTQ), a widely-used measure of trauma and early life adversity (Bernstein et al., 2003). We used CTQ Total Scores as a covariate in our primary and secondary analyses to identify variance in brain function and structure attributable to parenting style above and beyond that associated with childhood trauma, as the relationship between early life caregiving extremes and corticolimbic circuitry is well documented. Socioeconomic status (SES) was assessed using The MacArthur Scale of Subjective Social Status, which was developed to capture the common sense of social status across SES indicators by presenting a “social ladder” and asking individuals to place an "X" on the rung on which they feel they stand (Adler & Stewart, 2007). We used SES as a covariate in our primary and secondary analyses to isolate variance in corticolimbic circuit function and structure attributable to parenting style above and beyond that associated with socioeconomic status, as poverty can be interpreted as a chronic stressor and socioeconomic status has been shown to impact parenting style (Roubinov & Boyce, 2017).

### Amygdala Reactivity Task

Our experimental protocol consisted of four task blocks interleaved with five control blocks. The four task blocks consisted of one block each with fearful, angry, surprised, or neutral facial expressions presented in a pseudorandom order across participants. During task blocks, participants viewed a trio of faces and selected one of two faces (bottom) identical to a target face (top). Each task block consisted of six images, balanced for gender, all of which were derived from a standard set of pictures of facial affect (Ekman, 1976). During control blocks, participants viewed a trio of simple geometric shapes (circles and vertical and horizontal ellipses) and selected one of two shapes (bottom) identical to a target shape (top). Each control block consisted of six different shape trios. All blocks are preceded by a brief instruction (“Match Faces” or “Match Shapes”) that lasted 2 s. In the task blocks, each of the six face trios was presented for 4 s with a variable interstimulus interval (ISI) of 2-6 s (mean = 4 s) for a total block length of 48 s. A variable ISI was used to minimize expectancy effects and resulting habituation and maximize amygdala reactivity throughout the paradigm. In the control blocks, each of the six shape trios was presented for 4 s with a fixed ISI of 2 s for a total block length of 36 s. Total task time was 390 s.

### BOLD fMRI Data Acquisition

A semi-automated high-order shimming program was used to ensure global field homogeneity. A series of 34 interleaved axial functional slices aligned with the anterior commissure-posterior commissure plane were acquired using an inverse-spiral pulse sequence to reduce susceptibility artifacts (TR/TE/flip angle=2000 ms/30 ms/60; FOV=240mm; 3.75×3.75×4mm voxels; interslice skip=0). Four initial radiofrequency excitations were performed (and discarded) to achieve steady-state equilibrium. To allow for spatial registration of each participant’s data to a standard coordinate system, high-resolution three-dimensional structural images were acquired in 34 axial slices coplanar with the functional scans (TR/TE/flip angle=7.7 s/3.0 ms/12; voxel size=0.9×0.9×4mm; FOV=240mm, interslice skip=0).

### BOLD fMRI Data Pre-Processing

Anatomical images for each participant were skull-stripped, intensity-normalized, and nonlinearly warped to a study-specific average template in Montreal Neurological Institute (MNI) standard stereotactic space using ANTs (Klein et al., 2009). BOLD time-series for each participant were processed in AFNI (Cox, 1996). Images for each participant were despiked, slice-time-corrected, realigned to the first volume in the time series to correct for head motion, coregistered to the anatomical image using FSL’s Boundary Based Registration (Greve & Fischl, 2009), spatially normalized into MNI space using the non-linear warp from the anatomical image, resampled to 2mm isotropic voxels, and smoothed to minimize noise and residual difference in gyral anatomy with a Gaussian filter set at 6-mm full-width at half-maximum. All transformations were concatenated so that a single interpolation was performed. Voxel-wise signal intensities were scaled to yield a time series mean of 100 for each voxel. Volumes exceeding 0.5mm frame-wise displacement (FD) or 2.5 standardized DVARS (Nichols, 2017; Power et al., 2014) were censored from the subsequent GLM analyses.

### fMRI Quality Control

Quality control criteria for inclusion of a participant’s imaging data were: >5 volumes for each condition of interest retained after censoring for FD and DVARS and sufficient temporal signal-to-noise ratio (SNR) within the bilateral amygdala, defined as greater than 3 standard deviations below the mean of this value across participants. The amygdala was defined using a high-resolution template generated from 168 Human Connectome Project datasets (Tyszka & Pauli, 2016). Since we did not have *a priori* predictions regarding hemispheric differences, and to reduce the number of statistical tests, the extracted values were averaged across left and right hemispheres for further statistical analyses. Additionally, data were only included in further analyses if the participant demonstrated sufficient engagement with the task, defined as achieving at least 75% accuracy during the face matching condition.

### BOLD fMRI Data Analysis

Following preprocessing, the AFNI program 3dREMLfit (Cox, 1996) was used to fit general linear models for first-level fMRI data analyses. To obtain parameter estimates for each task block, we modeled only the respective block (convolved with the canonical hemodynamic response function) along with the adjacent half of the preceding and following control blocks, and a first order polynomial regressor to account for low frequency noise. This allowed for the estimation of the individual task block parameters while minimizing the influence of adjacent task blocks as well as low frequency noise across the entire run. Based on our prior work, the contrasts of interest for the current analyses were angry facial expressions > shapes and fearful facial expressions > shapes.

### Psychophysiological Interactions

Task-modulated functional connectivity was estimated using the generalized psychophysiological interaction (gPPI) toolbox (McLaren, Ries, Xu, & Johnson, 2012) in SPM12. Following preprocessing, deconvolved time courses averaged across the amygdala (anatomically defined) were extracted, and entered into first-level statistical models, which also included a psychological task regressor as well as an interaction term (seed*task). Individual beta images corresponding to the interaction term (seed*task) were then used in a second-level random effects model accounting for scan-to-scan and participant-to-participant variability to determine mPFC activation that varies as a function of amygdala reactivity and experimental condition. The mPFC mask was anatomically defined using SPM12 and included Brodmann Areas 10, 11, 12, 24, 25, and 32 using SPM12. Significance thresholds were set using an overall false detection probability based on 10,000 Monte Carlo simulations yielding a cluster-forming threshold of *p* < .001 and a cluster size of at least 98 contiguous voxels to achieve an overall family-wise error rate of α < .05.

### Diffusion Weighted Imaging

Following an ASSET calibration scan, two 2-min 50-s diffusion weighted imaging acquisitions were collected, providing full brain coverage with 2-mm isotropic resolution and 15 diffusion weighted directions (10-s repetition time, 84.9 ms echo time, b value 1,000 s/mm2, 240 mm field of view, 90° flip angle, 128×128 acquisition matrix, slice thickness=2 mm). Diffusion weighted images were processed according to the protocol developed by the Enhancing Neuro Imaging Genetics Through Meta-Analysis consortium (Jahanshad et al., 2013). In brief, raw diffusion weighted images underwent eddy current correction and linear registration to the non-diffusion weighted image in order to correct for head motion. These images were skull-stripped and diffusion tensor models were fit at each voxel using FMRIB’s Diffusion Toolbox (FDT; http://fsl.fmrib.ox.ac.uk/fsl/fslwiki/FDT) and the resulting two FA maps were linearly registered to each other and then averaged. Average FA images from all subjects were non-linearly registered to the ENIGMA-DTI target FA map, a minimal deformation target calculated across a large number of individuals (Jahanshad et al., 2013). The images were then processed using the tract-based spatial statistics (TBSS) analytic method (Smith et al., 2006) modified to project individual FA values onto the ENIGMA-DTI skeleton. Following the extraction of the skeletonized white matter and projection of individual FA values, left and right UF pathways of interest, defined using the Johns Hopkins University White Matter Tractography Atlas (Mori, Wakana, Nagae-Poetscher, & Van Zijl, 2006), were binarized to extract mean FA values each participant. Since we again did not have *a priori* predictions regarding hemispheric differences, and to reduce the number of statistical tests, the extracted FA values were averaged across left and right hemispheres for further statistical analyses (d’Arbeloff et al., 2018; Kim et al., 2019).

### Statistical Analyses

Mean individual contrast-related BOLD parameter estimates from functional clusters were entered into second-level analyses in SPSS, version 25 (IBM, Armonk, NY). To test our primary hypothesis, we ran a linear multiple regression analysis including all four PBI dimensions as predictor variables (maternal care, maternal overprotection, paternal care, paternal overprotection), extracted BOLD values for the contrast of angry facial expressions greater than shapes averaged across hemispheres as the sole outcome variable. We ran this regression first with no covariates, and then with age, sex, SES, and CTQ scores as covariates to explore the effects of PBI above and beyond effects of relevant confounding variables. To probe the specificity of any association, we conducted post hoc analyses by duplicating our initial model with outcome variable of extracted BOLD values for the contrast of fearful expressions greater than shapes.

After finding a significant association specifically between maternal overprotection and amygdala reactivity to angry facial expression, we focused subsequent analyses on this PBI dimension exclusively to reduce inflated false positives due to multiple tests. To this end, analyses testing our secondary hypotheses were conducted as simple, bivariate correlations between (1) maternal overprotection and extracted amygdala-mPFC gPPI values and (2) maternal overprotection and extracted FA values for the uncinate fasciculus. We first ran bivariate correlations with no covariates, then duplicated these models using partial correlation analyses in SPSS with age, sex, SES, and CTQ scores as covariates to explore the effect of maternal overprotection above and beyond effects of relevant confounding variables.

## RESULTS

### Participant Characteristics

Data were available from a maximum of 312 participants (170 women). Sample distributions and descriptive statistics for each PBI subscale score as well as SES and CTQ Total scores are detailed in Supplemental Table 1. The subsample (*n* = 168) of participants included in structural connectivity analyses did not significantly differ from the full sample (*N* = 312) on sex, age, race, CTQ Total, SES, or PBI subscale scores with the exception of maternal care (*t* = −2.40, *p* = 0.004). Zero-order correlations among all self-report measures are reported in Supplemental Table 2. There were no significant differences in PBI scores between participants who met criteria for one or more past or current DSM-IV diagnoses and those who did not (all *p*s > 0.5).

### Caregiving and Amygdala Reactivity

Consistent with our prior work, first-level analyses revealed robust bilateral amygdala reactivity to angry and fearful facial expressions across participants (Kim et al., 2018; Nikolova et al., 2014; Prather, Bogdan, & Hariri, 2013; Swartz et al., 2017). There were no significant differences in amygdala reactivity, amygdala-mPFC connectivity, or UF FA as a function of scanner (all *p*s > 0.2). Linear regression analyses using extracted BOLD parameter estimates from clusters exhibiting main effects of expression revealed a significant negative correlation between PBI maternal overprotection scores and amygdala reactivity to angry facial expressions (*Std. B* = 0.195, *p* = 0.009; Figure 1). There were no significant correlations between amygdala reactivity to angry facial expressions and paternal overprotection or paternal or maternal care (paternal overprotection: *Std. B* = −0.036, *p* = 0.613; maternal care: *Std. B* = 0.119, *p* = 0.091; paternal care: *Std. B* = −0.093, *p* = 0.167). The association between maternal overprotection and amygdala reactivity to angry facial expressions remained significant when controlling for age, sex, SES, and CTQ scores (*Std. B* = 0.181, *p* = 0.015). There were no significant correlations between any PBI subscales and amygdala reactivity to fearful facial expressions (maternal care: *Std. B* = 0.053, *p* = 0.460; maternal overprotection: *Std. B* = 0.073, *p* = 0.332; paternal care: *Std. B* = −0.034, *p* = 0.621; paternal overprotection: *Std. B* = −0.065, *p* = 0.365;); these associations remained non-significant when covarying for age, sex, SES, and CTQ scores. Given the specificity of this association to maternal overprotection, all subsequent analyses focused on this PBI scale to limit multiple comparisons. Full regression statistics for this primary analysis, including and excluding covariates, are reported in Supplemental Table 3.

**Figure 1.**
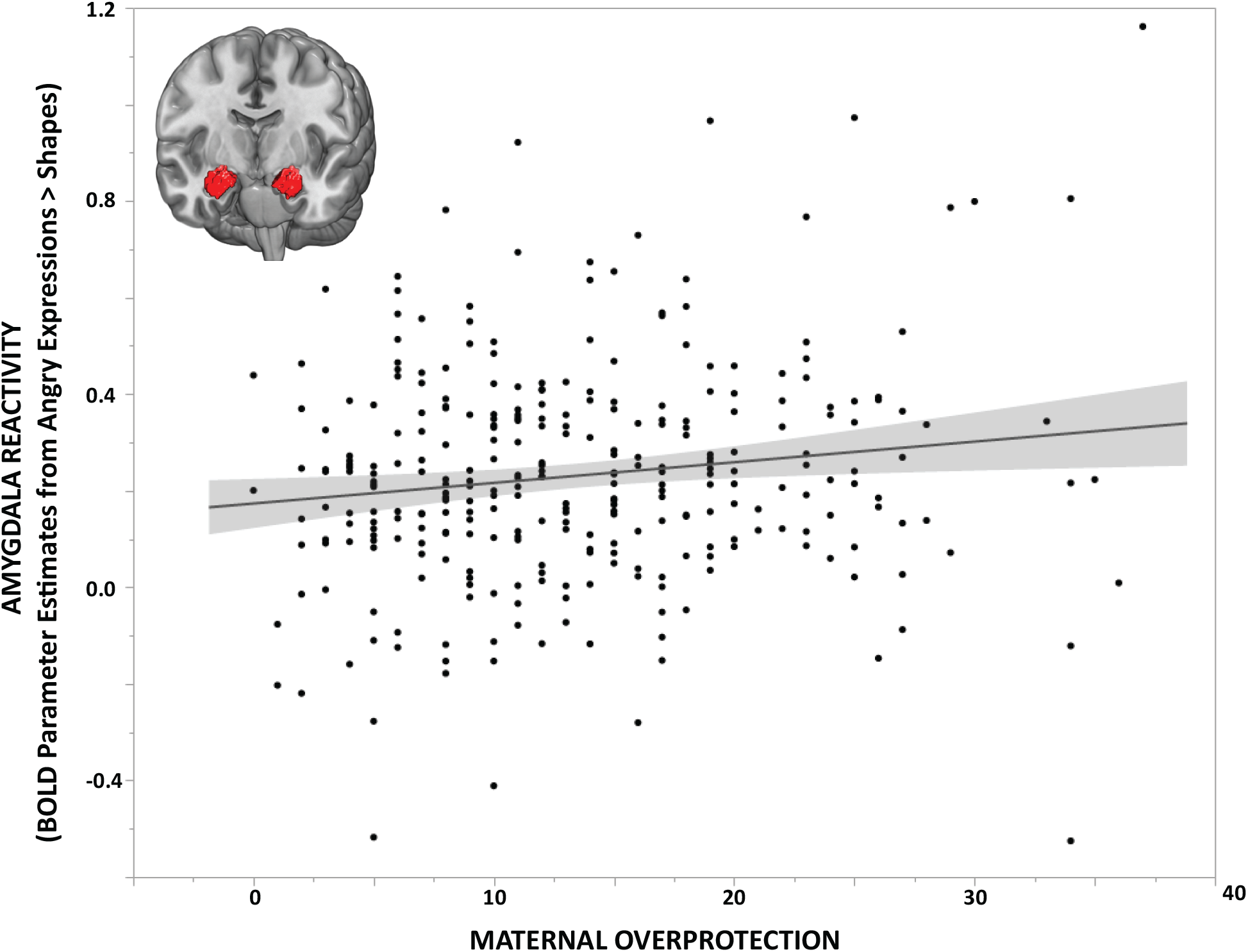
Maternal Overprotection and Amygdala Reactivity. Parental Bonding Instrument (PBI) maternal overprotection scores were positively associated with mean bilateral amygdala reactivity to interpersonal threat as indexed by extracted BOLD parameter estimates from clusters exhibiting main effects of angry facial expressions > shapes (*Std. B* = 0.195, *p* = 0.009; *N* = 312).

### Maternal Overprotection and Amygdala Functional Connectivity

Extending our primary finding, we next tested for an association between maternal overprotection and amygdala-mPFC functional connectivity when processing angry facial expressions. We extracted mean gPPI values across our mPFC mask generating a single value indicating the strength of task-modulated functional connectivity between the amygdala and mPFC for each participant. Bivariate correlation analysis in SPSS revealed no significant correlation between maternal overprotection and amygdala-mPFC functional connectivity (*r* = −0.009, *p* = 0.873). These correlations remained non-significant when controlling for age, sex, SES, and CTQ scores (*Partial r* = 0.029, *p* = 0.612).

### Maternal Overprotection and Amygdala Structural Connectivity

We next explored associations between maternal overprotection and amygdala-prefrontal structural connectivity. We extracted FA values such that each individual subject had a single value representing the white matter microstructural integrity of the UF, averaged across left and right hemispheres. Bivariate correlation analysis in SPSS revealed a significant negative correlation between maternal overprotection and UF FA (*r* = −0.166, *p* = 0.031; Figure 2). However, when controlling for age, sex, SES, and CTQ scores, this association was no longer significant (*Partial r* = −0.130, *p* = 0.097). In unpacking these correlations further, we found that the association between maternal overprotection and UF FA remained significant when controlling for sex (*Partial r* = −0.160, *p* = 0.039) or SES (*Partial r* = −0.162, *p* = 0.037), but was reduced to a trend level when controlling for age (*Partial r* = −0.148, *p* = 0.056) or CTQ (*Partial r* = −0.149, *p* = 0.055).

**Figure 2.**
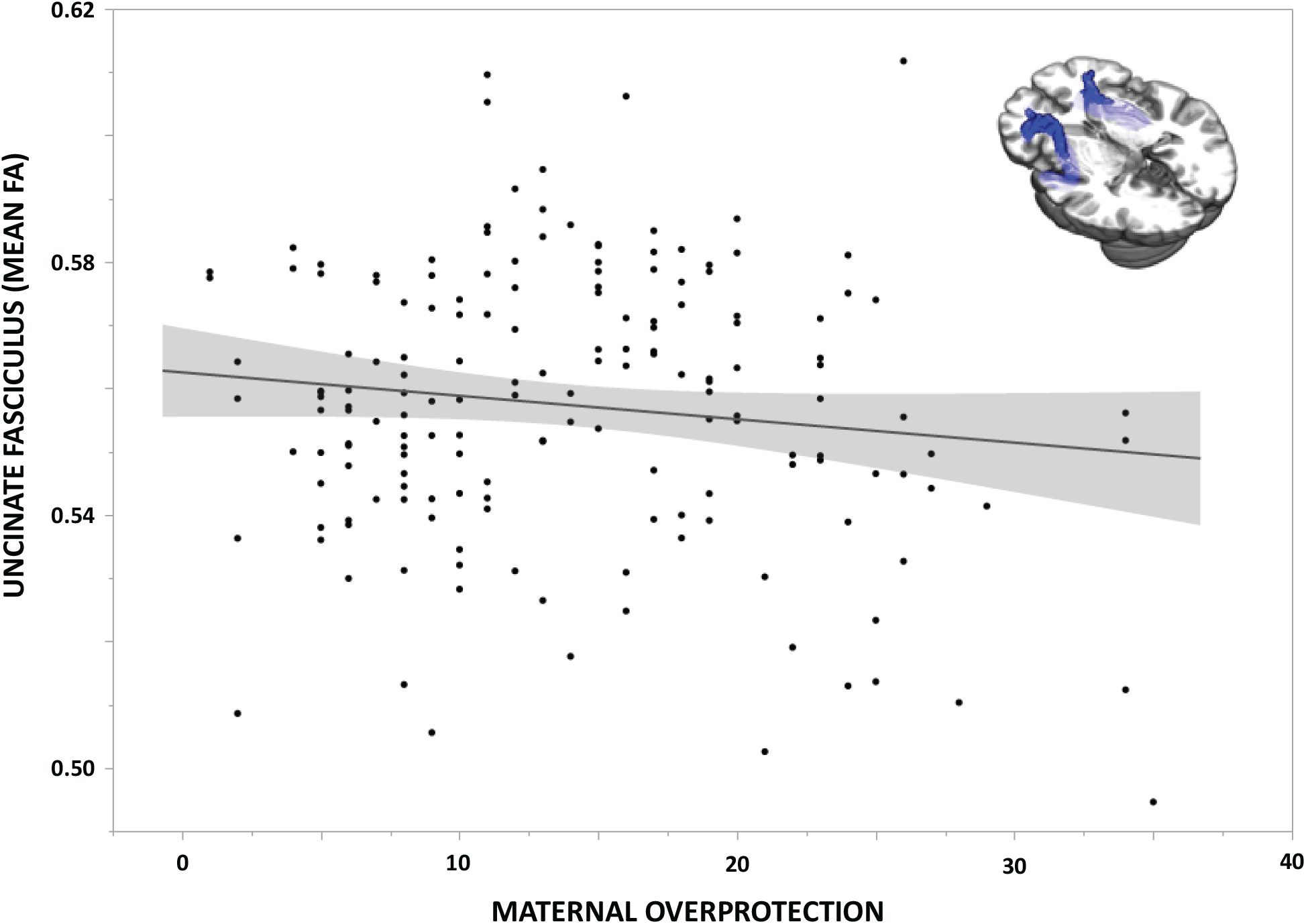
Maternal Overprotection and Amygdala Structural Connectivity. Paternal Bonding Inventory (PBI) maternal overprotection scores were negatively correlated with mean bilateral fractional anisotropy of the uncinate fasciculus (*r* = −0.166, *p* = 0.031; *n* = 168).

## DISCUSSION

The results of our study provide further evidence that normative variation in caregiving during childhood is associated with later behaviorally-relevant neural function and structure. First, while not a direct replication, we find support for our hypotheses regarding parental overprotection born of our previous work linking greater familial affective responsiveness with amygdala reactivity. We found that higher maternal overprotection but not paternal overprotection or maternal or paternal care is associated with increased amygdala reactivity to interpersonal threat in the form of angry facial expressions, above and beyond effects of potentially confounding variables such as age, sex, early life adversity, and socioeconomic status. Second, we were able to expand these findings into structural and functional connectivity between the amygdala and regulatory regions of the mPFC. Here we found that higher maternal overprotection is associated with decreased structural integrity of the uncinate fasciculus, a central white matter tract connecting the amygdala and mPFC. However, this correlation was non-significant when controlling for our variables of no interest, indicating that age and CTQ specifically absorb some of the variance captured by maternal overprotection on this phenotype. Contrary to our hypothesis, maternal overprotection was not significantly associated with functional connectivity between the amygdala and mPFC when processing angry facial expressions, regardless of covariates.

Our current findings are relevant for research on the buffering effects of parental presence. For example, maternal presence has been associated with suppression of amygdala reactivity in childhood but not adolescence (Gee et al., 2014). Thus, amygdala hyperactivity and weaker microstructural integrity of the uncinate fasciculus may reflect a childhood marked by overprotectiveness and, possibly, prolonged and therefore maladaptive maternal buffering. It is interesting to speculate that children who are relatively sheltered, particularly by their maternal caregiver, may not have sufficient opportunity to experience stress and subsequently fail to fully develop important structural pathways between the amygdala and mPFC. This impact on structure may then emerge as amygdala hyperactivity to interpersonal threat cues possibly in combination with insufficient prefrontal regulation. Such speculation, of course, cannot be directly tested in our cross-sectional data but requires longitudinal data ideally incorporating objective measurements of early upbringing and directly testing brain development over time.

Nevertheless, there is support for this speculation in both basic and clinical research. As discussed in our previous work (Farber et al., 2018), animal research illustrates that physiological hypersensitivity to threat, via behavioral freezing or amygdala reactivity, is most exacerbated when aversive stimuli are infrequent and unpredictable (Bouton, 2007; Fanselow & Tighe, 1988). It is possible that young adults who report higher overprotection by maternal caregivers may have experienced infrequent and unpredictable instances of interpersonal threat during childhood and, as such, exhibit amygdala hyperreactivity to signals of such threat (i.e., angry facial expressions) paired with relatively weakened white matter structural integrity connecting limbic to prefrontal regions.

Our work, of course, is not without limitations, the most pressing of which is that our findings are correlational and derived from cross-sectional data. To establish directional effects between parenting style and brain function and structure requires multiple timepoints of each construct within a longitudinal study design. At present, we can merely speculate in our interpretation of directionality of the observed associations. Similarly, these analyses are correlational in nature. We did not have data available to explore the potentially bidirectional nature of parenting style wherein it is possible that child temperament and parent temperament interact to influence parent behavior, which then influences brain development in the child. In addition, we relied on self-report measures of parenting; therefore, our findings may be subject to reporting bias and are likely more representative of the perception of events rather than of objective events. Also of note, our participants could be described as W.E.I.R.D — Western, Educated, Industrialized, Rich, and Democratic (Henrich, Heine, & Norenzayan, 2010). The large majority of published studies utilize such samples, but it is nevertheless important to note that the present study sample is not population representative and that this limits the generalizability and associated external validity of our results. Future studies in more diverse and ideally population representative samples are necessary to evaluate the extent to which our current findings are relevant more broadly. Likewise, future work should examine these patterns in more diverse family structures. While we limited our present analyses to adolescents raised in two-parent, one maternal and one paternal caregiver households, it is important to explore effects of normative caregiving in individuals raised in single-parent households, two-parent same-sex households, and other familial configurations.

Limitations exist with our fMRI task as well. We focused on amygdala reactivity from contrasts of emotional facial expressions against our control shape matching condition rather than direct contrasts of different emotional expressions as the latter did not yield statistically significant results even with the power afforded by our relatively large sample. Nevertheless, claims of expression-specific effects as a function of maternal overprotection would be strengthened by testing differential amygdala reactivity to angry versus fearful expressions. In addition, our fMRI task precludes examination of amygdala-mPFC functional connectivity during explicit emotion regulation. Thus, we cannot provide functional results in parallel with our structural results. Lastly, a growing number of studies report poor test-retest reliability of amygdala reactivity during tasks using emotional facial expressions as stimuli, including the task used in our protocol (Lipp, Murphy, Wise, & Caseras, 2014; Lois, Kirsch, Sandner, Plichta, & Wessa, 2018; Nord, Gray, Charpentier, Robinson, & Roiser, 2017; Plichta et al., 2012; Sauder, Hajcak, Angstadt, & Phan, 2013). Thus, task-elicited functional activation in *a priori* regions of interest may not be well suited for individual differences research. That said, the current associations are remarkably consistent with our earlier associations between amygdala reactivity and normative variability in caregiving.

These limitations notwithstanding, our current findings further extend the literature on the brain effects of caregiving extremes to more subtle, normative variability. Our findings suggest that overprotective maternal caregiving is associated with increased amygdala reactivity to explicit signals of interpersonal threat and decreased microstructural integrity of a pathway between the amygdala and mPFC known to support emotion integration and regulation (Lee, Heller, Van Reekum, Nelson, & Davidson, 2012) and resilience to mood and anxiety disorders (Etkin & Wager, 2007; Koenigs & Grafman, 2009; Murray, Wise, & Drevets, 2011; Tottenham, 2018). It is critical that we continue to build on the foundational work by early adversity researchers and investigate not if, but how normal range parenting is associated with brain phenotypes underlying risk and resilience for psychopathology. In the meantime, to borrow from research on the importance of risky play for children, our work suggests it may be ideal for caregivers to keep children “as safe as necessary,” not “as safe as possible” (Brussoni, Olsen, Pike, & Sleet, 2012).

## ACKNOWLEDGMENTS

We thank the Duke Neurogenetics Study participants as well as the staff of the Laboratory of NeuroGenetics. The Duke Neurogenetics Study was supported by Duke University and NIH grants R01DA031579 and R01DA033369. ARH is further supported by NIH grant R01AG049789. The Duke Brain Imaging and Analysis Center’s computing cluster, upon which all DNS analyses heavily rely, was supported by the Office of the Director, National Institutes of Health under Award Number S10 OD 021480.

