## Supplemental Tables 1-3 for "Maternal Overprotection in Childhood is Associated with Amygdala Reactivity and Structural Connectivity in Adulthood" (from another version of this preprint: version 1, 2019-02-01)

**Table 1.** Sample descriptive statistics: Parental Bonding Instrument (PBI) subscales, Childhood Trauma Questionnaire (CTQ) total scores, and SES as indexed by Social Ladder rating.

|  | *Range* | *Mean* | *SD* | *Skewness* | *Kurtosis* |
| --- | --- | --- | --- | --- | --- |
| **Maternal Care** | 5 – 35 | 27.77 | 5.47 | -1.15 | 0.83 |
| **Maternal Protection** | 0 – 37 | 13.41 | 7.63 | 0.69 | 0.07 |
| **Paternal Care** | 4 – 34 | 24.89 | 6.79 | -0.86 | 0.22 |
| **Paternal Protection** | 0 – 37 | 10.11 | 7.22 | 0.82 | 0.46 |
| **CTQ** | 25 – 75 | 32.80 | 8.01 | 1.99 | 5.51 |
| **SES** | 2 - 11 | 7.88 | 1.73 | -0.76 | 0.52 |

**Table 2.** Zero-order (Pearson) correlations: PBI subscales, Childhood Trauma Questionnaire (CTQ) total score, and Socioeconomic Status (SES) as indexed by Social Ladder rating.

|  | **M Care** | **M Prot** | **P Care** | **P Prot** | **CTQ** | **SES** |
| --- | --- | --- | --- | --- | --- | --- |
| **Maternal Care** | *1.00* | -0.457 ** | 0.495 ** | -0.267 ** | -0.550 ** | 0.122 * |
| **Maternal Protection** | -0.457 ** | *1.00* | -0.303 ** | 0.568 ** | 0.347 ** | -0.100 |
| **Paternal Care** | 0.495 ** | -0.303 ** | *1.00* | -0.359 ** | -0.474 ** | 0.142 * |
| **Paternal Protection** | -0.267 ** | 0.568 ** | -0.359 ** | *1.00* | 0.342 ** | -0.216 ** |
| **CTQ** | -0.550 ** | 0.347 ** | -0.474 | 0.342 ** | *1.00* | -0.268 ** |
| **SES** | 0.122 * | -0.100 | 0.142 * | -0.216 ** | -0.268 ** | *1.00* |

** *p* < 0.001

* *p* < 0.05

**Table 3.** Regression statistics for our primary multiple regression analysis examining all Parental Bonding Inventory (PBI) scales (maternal care, paternal care, maternal protection, paternal protection) and amygdala reactivity (a) without covariates and (b) including age, sex, Childhood Trauma Questionnaire (CTQ) total score, and Socioeconomic Status (SES) as indexed by Social Ladder rating.

**(a)** Model Summary: *R* = 0.182, *R^2^* = 0.033, *Adjusted R^2^* = 0.020

| ANOVA | | | | | |
| --- | --- | --- | --- | --- | --- |
|  | *Sum of Squares* | *df* | *Mean Square* | *F* | *p* |
| **Regression** | 0.531 | 4 | 0.133 | 2.607 | 0.036 |
| **Residual** | 15.57 | 306 | 0.051 |  |  |
| **Total** | 16.101 | 310 |  |  |  |

| Coefficients | | |
| --- | --- | --- |
|  | *Std. B* | *p* |
| **Maternal Care** | 0.119 | 0.091 |
| **Maternal Protection** | 0.195 | 0.009 |
| **Paternal Care** | -0.093 | 0.167 |
| **Paternal Protection** | -0.036 | 0.613 |

**(b)** Model Summary: *R* = 0.225, *R^2^* = 0.051, *Adjusted R^2^* = 0.026

| ANOVA | | | | | |
| --- | --- | --- | --- | --- | --- |
|  | *Sum of Squares* | *df* | *Mean Square* | *F* | *p* |
| **Regression** | 0.816 | 8 | 0.102 | 2.015 | 0.044 |
| **Residual** | 15.285 | 302 | 0.051 |  |  |
| **Total** | 16.101 | 310 |  |  |  |

| Coefficients | | |
| --- | --- | --- |
|  | *Std. B* | *p* |
| **Sex** | -0.038 | 0.500 |
| **Age** | -0.031 | 0.588 |
| **SES** | 0.125 | 0.035 |
| **CTQ** | 0.079 | 0.276 |
| **Maternal Care** | 0.137 | 0.071 |
| **Maternal Protection** | 0.181 | 0.015 |
| **Paternal Care** | -0.075 | 0.279 |
| **Paternal Protection** | -0.015 | 0.841 |
